# Physical Obstacles Constrain Behavioral Parameter Space of Successful Localization in Honey Bee Swarms

**DOI:** 10.1101/2022.05.17.492366

**Authors:** Dieu My T. Nguyen, Michael L. Iuzzolino, Orit Peleg

**Affiliations:** Department of Computer Science, University of Colorado Boulder, Boulder, Colorado, USA; BioFrontiers Institute, University of Colorado Boulder, Boulder, Colorado, USA; Santa Fe Institute, Santa Fe, New Mexico, USA

## Abstract

Honey bees (*Apis mellifera* L.) localize the queen and aggregate into a swarm by forming a collective scenting network to directionally propagate volatile pheromone signals. Previous experiments show the robustness of this communication strategy in the presence of physical obstacles that partially block pheromone flow and the path to the queen. Specifically, there is a delay in the formation of the scenting network and aggregation compared to a simple environment without perturbations. To better understand the effect of obstacles beyond temporal dynamics, we use the experimental results as inspiration to explore how the behavioral parameter space of collective scenting responds to obstacle. We extend an agent-based model previously developed for a simple environment to account for the presence of physical obstacles. We study how individual agents with simple behavioral rules for scenting and following concentration gradients can give rise to collective localization and swarming. We show that the bees are capable of navigating the more complex environment with a physical obstacle to localize the queen and aggregate around her, but their range of behavioral parameters is more limited and less flexible as a result of the spatial density heterogeneity in the bees imposed by the obstacle.

## Introduction

Social insect groups often navigate complex and unknown environments. To do so, group members must effectively communicate. Insects, such as honey bees, often exchange information and coordinate group processes by communicating via pheromones, volatile chemical signals that decay rapidly in time and space (Conte and Hefetz, 2008; Lensky and Cassier, 1995). In the context of honey bee swarm formation around the queen, worker bees localize the queen by following her pheromones and propagate the signals about her location by “scenting” (McIndoo, 1914; Peters et al., 2017; Nguyen et al., 2021b). The scenting behavior consists of a given bee sensing local pheromone concentration above a given threshold and releasing pheromones from the Nasonov gland while rigorously fanning its wings to disperse the signals to other bees. Wing fanning creates a directional bias in the flow of pheromones, allowing bees farther away from the queen to sense the signal and further propagate them. This collective scenting strategy creates an effective communication network for localization and aggregation in honey bee swarms (Nguyen et al., 2021b).

Nguyen et al. (2021a) experimentally showed the robustness of collective scenting in the presence of obstacles that partially block pheromone flow and the open path to the queen. Compared to a simple environment without any obstacle, the more complex environment requires more time to explore and navigate. However, the bees still effectively employ the scenting strategy to overcome the obstacle and aggregate around the queen (Fig. 1B).

**Figure 1:**
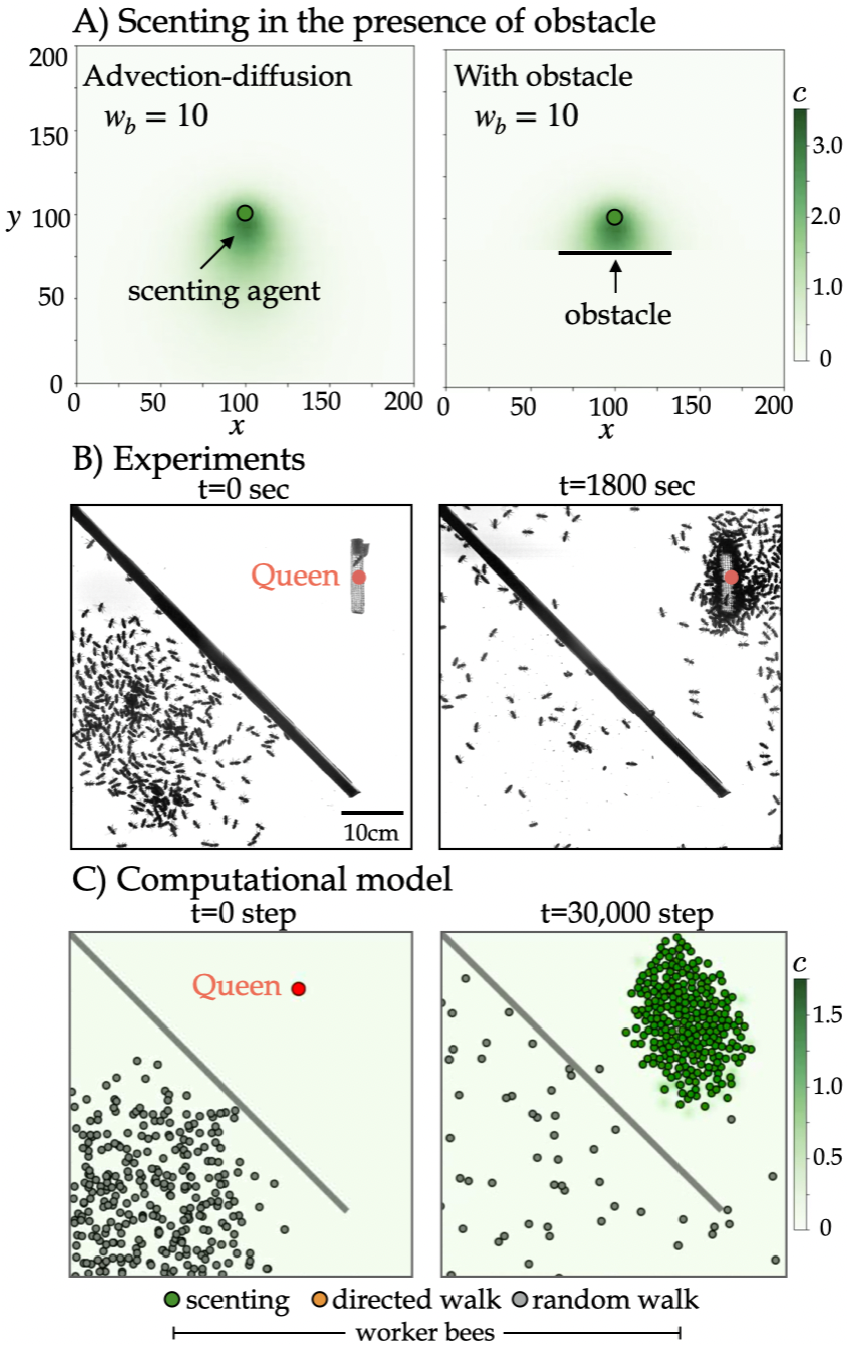
Experiments and model of collective scenting in the presence of a physical obstacle. A) A scenting bee agent can produce a pheromone signal with directional bias with advection-diffusion (*w_b_* = 10). A physical obstacle can partially block the signal. B) An experiment where worker bees use scenting to navigate the obstacle and localize the queen. C) A computational model simulating pheromone diffusion and honey bee scenting behavior in presence of obstacle.

Inspired by the experiments, we turned to computational modelling to further explore how physical obstacles affect the parameter space that dictates the dynamics of localization and aggregation based on collective scenting. A previous work modeling social amoeba aggregation by chemical signaling has shown the system’s robustness to physical obstacles, with agents releasing isotropic chemicals that diffuse axi-symmetrically (Fatès, 2010). To model honey bee chemical signaling, a previous study (Nguyen et al., 2021b) used agent-based modeling to study how simple behavioral rules can generate complex collective behaviors and took into consideration the directional bias of signals that provide directional information to group members. The authors studied the effect of two behavioral parameters that the bees may vary with input from the environment—the directional bias representing the magnitude of wing-fanning and the concentration threshold above which they can detect the signal. The model showed the importance of directional signals seen in the scenting strategy in efficient localization and aggregation around the queen that can avoid less desired outcomes, such as small clusters far from the queen. In this study, we build upon this model by adding physical obstructions to the system (Fig. 1A,C). Per the experiments in Nguyen et al. (2021a), we expect to observe the robustness of the bee communication system in the more complex environment. More importantly, by modeling, we aim to gain insights into how the behavioral parameter space responds to the presence of obstacle. The insights may contribute to designs and improvements in non-biological systems with individuals that are limited to local interactions but must coordinate group processes in complex environments, such as robots that must navigate obstructions (Abiyev et al., 2010).

## Methods

### Experimental setup & analysis

We followed methods described in detail in Nguyen et al. (2021b) for experiments without obstacle and in Nguyen et al. (2021a) with obstacle. As this paper mainly focuses on modeling, we briefly summarize the methods. The backlit arena (50×50×1.5 cm) is semi-2-D to prevent flying, as bees have been shown to scent while standing (McIndoo, 1914). The experiments are recorded aerially with a video camera (4k resolution, 30 fps). The queen is kept in a cage (10.5×2.2×2.2 cm) at the top right corner (Fig. 1B). A wooden bar is placed diagonally for the obstacle condition. Workers are placed at the bottom left corner, and a plexiglass sheet encloses the arena. For each environmental condition, we report results for three experiments with similar number of worker bees ranging from 240 to 380. Five experiments per condition were presented in Nguyen et al. (2021a); we present three per condition here for consistency in number of bees in the simulation (*N* = 300).

To automatically detect scenting bees and their orientations, we use computer vision approaches presented in Nguyen et al. (2021b) (Appendix Fig. A1). We detect individual bees (i.e., *x, y* centroids) by Otsu’s method of adaptive thresholding, and morphological transformations (Otsu, 1979; Dougherty, 1992). To classify a bee as scenting or non-scenting, we train a ResNet-18 convolutional neural network (CNN) model (He et al., 2016) that achieves 95.17% test accuracy. We create a regression model for orientation estimation, which achieves 96.71% test accuracy.

We then reconstruct attractive surfaces to correlate the scenting events with the spatiotemporal density of bees. For each scenting bee *i* at time *t*, we define its position as 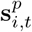, and its direction of scenting as 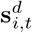 (unit vector). Assuming the scenting bees provide directional information to non-scenting bees, we treat 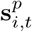 and 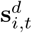 as a set of gradients that define a minimal surface of height *f*(*x, y, t*). Thus, *f*(*x, y, t*) corresponds to the probability that a randomly moving non-scenting bee will end up at position (*x, y*) by following the scenting directions of scenting bees: *f*(*x, y*) = ∑_∀∇*f*_ ∫∇*dxdy* where 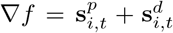. We regularize the least squares solution of surface reconstruction from its gradient field, using Tikhonov regularization (Harker and O’Leary, 2008, 2011).

Finally, we obtain some time-series properties. The number of scenting bees over time is presented as a rolling mean with the window size of 100 frames. The average distance to the queen is computed as the average distance of all black pixels to the queen’s location, as the bee detection method cannot detect every single individual bee when they touch or overlap. The queen’s cage and the obstacle are stationary, thus the remaining black pixels in the arena represent only the moving bees and allow us to use this proxy. For each property, we average the time-series data across all experiments for each condition and obtain the standard deviation.

### Modeling pheromone diffusion

We model pheromone advection-diffusion using the 2-D diffusion partial differential equation to describe pheromone concentration, *C*(*x, y, t*), at a position and time:

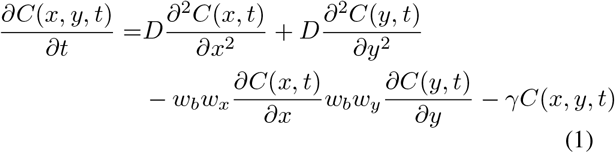

where *C*(*x, y, t*) is the concentration at position [*x, y*] at time *t*, *w_x_* and *w_y_* are the *x* and *y* components of emission vector respectively, *D* is the diffusion coefficient, and *γ* is the decay constant. The behavioral parameter representing the directional bias, *w_b_*, is the magnitude of the advection–diffusion of pheromone released by a bee (Fig. 1A). Treating a single scenting bee as a point source of localized and instantaneous pheromone emission, we solve Eq. 1:

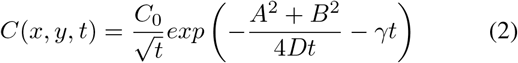

where *C*_0_ is the initial concentration, *A* = (*x* − *w_b_w_x_t*), and *B* = (*y* − *w_b_w_y_t*). The environmental parameters of the model include: the size of the 2–D arena (*X_min_* and *X_max_*) and the size of a grid cell (*δ_X_*), the start and final time of the simulation (*t_i_* and *t_f_*) and the time integration constant (*δ_t_*).

Similar to the experiments, we model the physical obstacle as a diagonal linear bar of pixels with a small opening. Pheromones cannot diffuse past the obstacle by a line-of-sight method. The obstacle forms a line segment 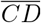, and segment 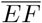 forms between the scenting agent and a given pixel. Intersection of the lines indicates that the pheromone from the scenting agent does not reach the pixel. To find the point of intersection, we solve the matrix equation: 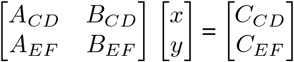 where *A_CD_* is the slope of 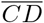, *A_EF_* is the slope of 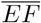, *B_CD_* = −1, *B_EF_* = −1, *C_CD_* is the negative y-intercept of 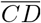, and *C_EF_* is the negative y-intercept of 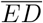. If a solution exists, we check if the intersection point lies on both lines. If there is no solution or the solution does not lie on both lines, there is no intersection and the pheromone from the source at *E* is present in the pixel at *F*.

### Modeling behavioral rules

In a discrete 2-D arena (Appendix Fig. A2A), the queen is stationary and frequently releases pheromone isotropically, i.e., without directional bias (*wb* = 0). She is the global point of convergence for the swarm. The behavioral rules of workers are: (1) A worker bee performs a random walk. Based on her distance to the queen, the bee detects the queen’s pheromone if above the threshold (*T*). (2) If *T* is met, the bee orients towards the direction up the gradient. The negative vector of the gradient scaled by *w*_b_ is the direction to emit pheromone and disperse it via wing-fanning. The bee then either walks up the gradient (Appendix Fig. A2B) or stands still for a certain time to emit and fan her own pheromones, each event with a 0.5 probability. (3) Bees that detect this cascade of secondary signals will follow the same rules to head towards maximum pheromone concentration or scent and further propagate the information.

We formalize the worker bees’ behavior as a probabilistic state machine (PSM) (Rabin, 1963). The PSM consists of a set of finite states that describe bee behavior and a probabilistic transition matrix for how a bee may change from one state to another. Specifically, the state model *SM_worker_* = (*S, s*_0_, *I, M*), associated with each worker, defines her set of behavioral rules within the environment, and *SM_worker_* components are fixed across all worker bees: *S* = {*randomWalk, directedWalk, thresholdMet, emit, fan*} is a set of finite states, where the variable *randomWalk* is a random walk when the threshold is not met, *directedWalk* is the walk up the concentration gradient, *thresholdMet* is when the threshold is met, *emit* is the instantaneous release of pheromone, and *fan* is the wing fanning at a constant position. *s*_0_ = *randomWalk* is the initial state of each bee. *I* = {*t_i_, c_i_*}, is a set of flags for the input conditions on state transitions, where for a given bee, *t_i_* is a counter for the time that bee is in the *fan* state and *c_i_* is the concentration at that bee’s position.

For the transition matrix *M*, there are two relevant parameters, *P_w_* and *T*, representing the emission period made of the *emit* and the *fan* state and the threshold over which a bee can be activated from state *randomWalk*. Table 1 provides the conditions and probabilities for transitioning from the current state, *s_c_*, to the next state, *s_n_*.

**Table 1:** Probabilistic state machine transition matrix for honey bee behavioral rules. Variables *randomWalk, thresholdMet*, and *directedWalk* abbreviated as *rWalk, tMet*, and *dWalk*, respectively.

| $s_c \backslash s_n$ | rWalk | tMet | emit | fan | dWalk |
| --- | --- | --- | --- | --- | --- |
| rWalk | $c_i < T$ | $c_i \geq T$ | 0 | 0 | 0 |
| dWalk | $c_i < T$ | $c_i \geq T$ | 0 | 0 | 0 |
| tMet | 0 | 0 | 0.5 | 0 | 0.5 |
| emit | 0 | 0 | 0 | 1 | 0 |
| fan | $t_i \geq P_w \wedge c_i < T$ | $t_i \geq P_w \wedge c_i \geq T$ | 0 | $t_i < P_w$ | 0 |

We compute the gradient of pheromone concentration for a given bee to find the direction of greatest local change:

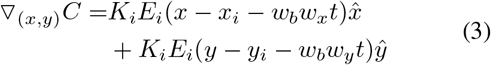

where 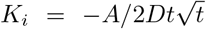 and *E_i_* = *exp*(−(*x* − *x_i_* − *w_b_w_x_t*)^2^ + (*y* − *y_i_* − *w_b_w_y_t*)^2^/4*Dt* − *γt*), *x, y* are the position of the activated bees, and *x_i_, y_i_* are the position of the scenting bees (i.e., pheromone source) *i*. The cumulative gradient for the concentration at a single bee’s position is the sum of the normalized gradients resulting from each pheromone source or emitting bee *i*:

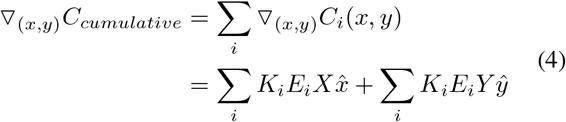

where *X* = *x* − *x_i_* − *w_b_w_x_t* and *Y* = *y* − *y_i_* − *w_b_w_y_t*. This gradient defines the vector that points in the direction of the bee’s heading for its directed walk. The negative vector of the gradient is the direction for this bee’s pheromone emission for signal propagation, and thus its *x* and *y* components make up the *w_x_* and *w_y_* terms of Eq. 2.

Each discrete pixel or cell contains only one bee at a time. Pixels that make up the physical obstacle do not contain any bees. Upon its next movement, a bee agent checks if the intended pixel is already occupied by another bee or the obstacle; if so, the bee chooses another nearby pixel within 45°over five iterations until it stays at the current position. Based on the model parameterization in Nguyen et al. (2021b), we explore a range of values for the behavioral parameters, the directional bias *w_b_* and concentration threshold *T*, that bees could adjust based on input from the environment. For each combination of the parameters, we repeat the simulation three times per condition. Other parameters remain constant across all simulations (Appendix Table 2). The algorithm is presented in Appendix Algo. 1.

### Construction of phase diagrams

We extract several properties from the simulation data for time-series analyses: the bees’ average distance to the queen, the average number of scenting bees, and the average distance to the queen from the farthest active bee. To characterize the growth of the queen’s cluster size and the number of clusters that form, we use the density-based spatial clustering of applications with noise (DBSCAN) algorithm (*ε*: 0.25, minimum number of bees to form a cluster: 5) to cluster bees at every time step (Ester et al., 1996).

To characterize the collective scenting behavior determined by the behavioral parameters (*w_b_, T*), we define four possible phases (i.e., the outcome of simulations, rather than a period in a time sequence) as previously seen in Nguyen et al. (2021b): phase 1 of small clusters of bees spread throughout the arena, phase 2 of bees reaching the queen’s vicinity by random walk, phase 3 of bees swarming around the queen by forming a percolation network of senders and receivers of pheromone signals created by scenting bees as seen in the experiments, and phase 4 of no clustering at all. To construct the phase diagrams, we use three properties to distinguish the phases: the final number of clusters, the final queen’s cluster size, and the distance of the farthest active bee to the queen. We sequentially applied the following conditions to each simulation identify its phase group: 1) If the final number of clusters > 1.5: Phase 1 with many small clusters; 2) If the final number of clusters 0 - 1.5 and the final queen’s cluster size < 250 bees: Phase 4 with no clustering; 3) If the farthest active distance < 4.0: Phase 2 with clustering at the queen’s location via random walk; 4) If the farthest active distance ≥ 4.0: Phase 3 with clustering at the queen’s location via a scenting percolation network.

## Results

### Bees navigate obstacles by collective scenting

As previously presented in Nguyen et al. (2021a), the experiments comparing the localization and aggregation dynamics in the presence and absence of physical obstacles show that bees are able to solve the problem in both conditions by employing the collective scenting strategy. We show snapshots of the experiment and the corresponding attractive surfaces for an example experiment without obstacles (*N* = 320) in Fig. 2A. Over 1800 seconds or 30 minutes, the bees activate a scenting network early (around *t* = 140 sec), as reflected in the surface in which the scenting directions collectively point to the queen’s area (i.e., surface regions of higher f values). Most of the bees have formed a swarm around the queen around *t* = 900 sec or 15 min. In the presence of obstacles, the bees generally require more time and exploration to form the scenting network. In Fig. 2B, we show snapshots of an example obstacle experiment (N = 310), in which bees search around the space behind the bar until a few bees find the opening and begin forming the collective scenting network at around 900 sec or 15 min. Most bees swarm around the queen by *t* = 1800 sec (30 min).

**Figure 2:**
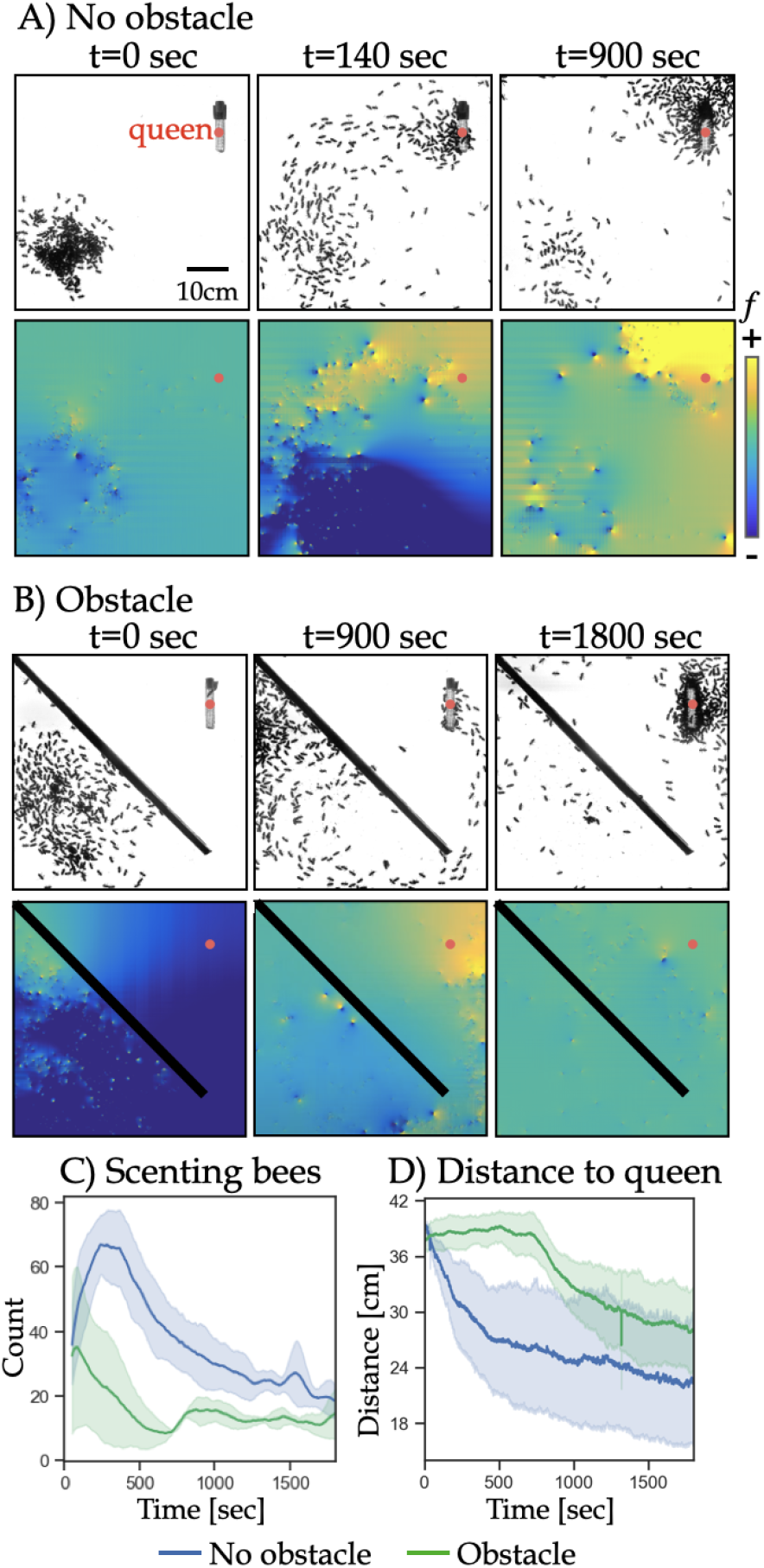
Experiments. A) Snapshots of an experiment where worker bees are in a semi-2D arena with a caged queen without obstacles. The corresponding attractive surfaces *f* show the scenting events correlating to the spatial-temporal density of bees. B) Snapshots and surfaces of an experiment where bees are initially placed on on side of a bar obstacle and the queen is on the other side. C) The average number of scenting bees over time for both conditions. D) The average distance to the queen over time.

To quantitatively compare the aggregation process over time for the two conditions, we analyze the number of scenting bees over time (averaged over three experiments for each condition, with shaded area showing the standard deviation) in Fig. 2C. Without the obstacle, there is a sharp peak in the early phase of the experiment (around 350 sec) when bees quickly form the scenting network and a gradual decrease as most bees have clustered around the queen. In the presence of the obstacle, there is also a very early peak (around 50 sec); however, this peak of scenting occurs behind the obstacle before the bees find the opening. A smaller peak around 900 sec occurs when bees find the opening and forms a scenting network where the attractive surface shows the scenting directions oriented towards the queen (Fig. 2B). Further, we compared the average distance to the queen over time (Fig. 2D). The distance sharply decreases early in the absence of an obstacle. With an obstacle, the plateau from the start of the experiment until approximately 800 sec indicates the time the bees spend behind the obstacle until the first bees explore and find the opening.

### Model shows constraints in behavioral parameter space in the presence of obstacles

The experimental results indicate the robustness of the collective scenting strategy in the presence of a physical obstacle and provide insights into the temporal dynamics of the aggregation process in different environments. The experiments are an inspiration for the agent-based model where we further explore the behavioral parameters behind the mechanisms of the collective scenting and localization process. With and without physical obstacles, the model shows the four distinct phases defined in the Methods section:

Phase 1: Low values of both *w_b_* and *T* produce small clusters of bees (Fig. 3A). Without an obstacle, signals reach the entire swarm and lead to clusters earlier than with an obstacle (around 900 and 3000 time steps, respectively).
Phase 2: High values of *T* produce swarms around the queen only via the bees that slowly reach the queen by random walk (Fig. 3B). When the obstacle is present, more time is required for bees to spread out throughout the arena (after 1500 time steps compared to by 900 time steps).
Phase 3: High values of *w_b_* and low values of *T* lead to the percolation network of scenting (Fig. 3C). Without the obstacle, the network has begun to form by around 900 time steps, while it takes around 5000 time steps with the obstacle. Although bees in phases 2 and 3 eventually cluster at the queen’s location, pheromone signals reach a much farther distance in phase 3 than in phase 2.
Phase 4: Very high values of *T* and *w_b_* lead to no worker bees ever activated to scent or perform the directed walk up the gradient, and therefore no clustering (Fig. 3D).

**Figure 3:**
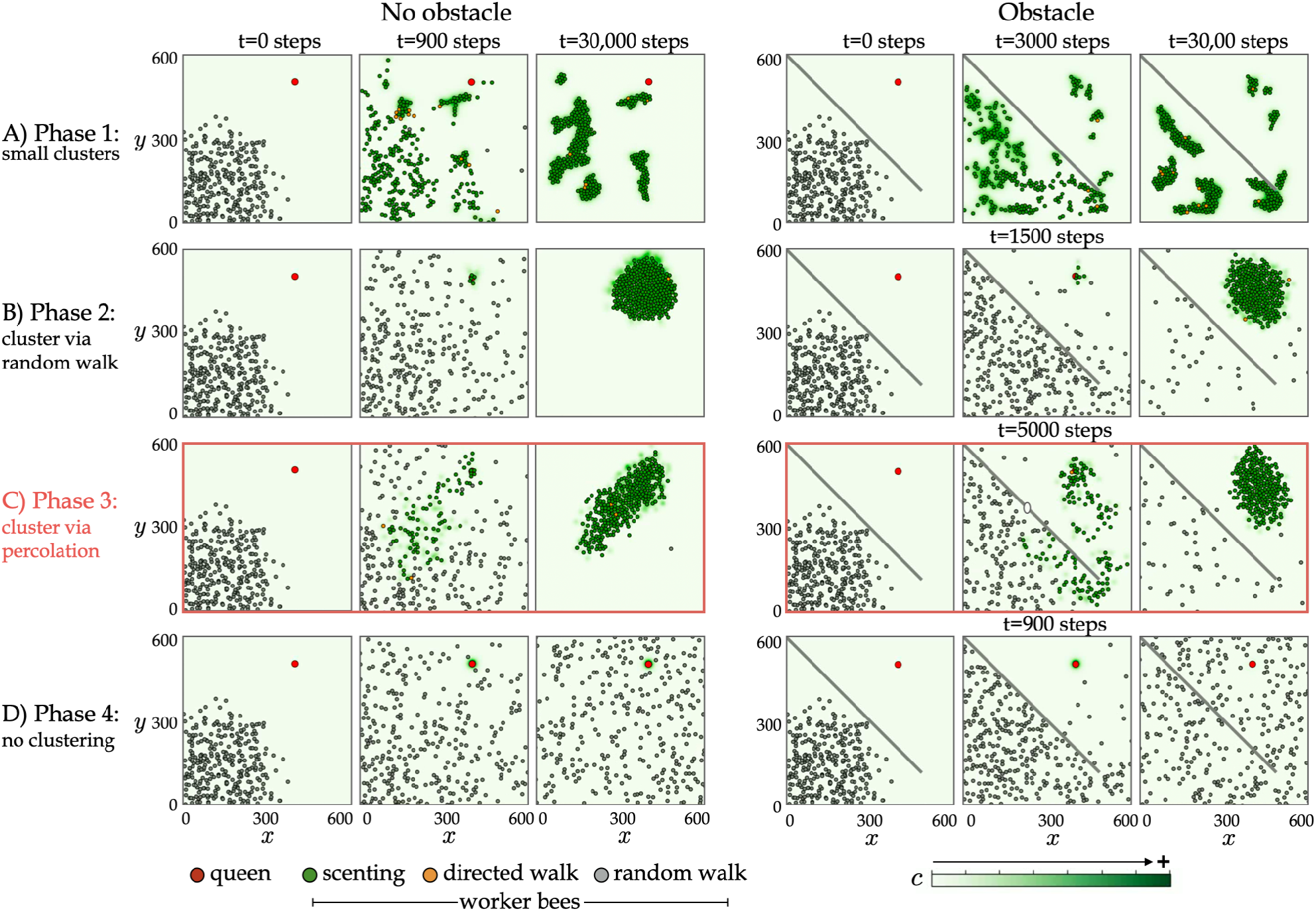
Simulations of four different phases. The queen is a red circle at the top right corner. Worker bees are circles colored by their internal state: scenting (green), performing a directed walk up the gradient (orange), and performing a random walk (gray). The instantaneous pheromone concentration *C*(*x, y, t*) corresponds to the green color scale. Common simulation parameters are *N* = 300, *C*_0_ = 0.0575, *D* = 0.6, and *γ* = 108. A) Phase 1 where bees aggregate into small clusters; *w_b_* = 0, *T* = 0.0001 for both conditions. B) Phase 2 where bees cluster around the queen via random walk; *w_b_* = 30, *T* = 0.5 for no obstacle; *w_b_* = 40, *T* = 0.2 for obstacle. C) Phase 3 where bees create a percolating network of senders and receivers of the pheromone signal to cluster around the queen; *w_b_* = 50, *T* = 0.025 for no obstacle; *w_b_* = 50, *T* = 0.075 for obstacle. D) Phase 4 where no clustering occurs; *w_b_* =60, *T* =1.0 for both.

Although all four phases are present in both conditions, the presence of the physical obstacle affects the phase areas and boundaries. The phase diagram of four phases as determined by (*w_b_, T*) for simulations without the obstacle is shown in Fig. 4A. Treating the phase diagram as an image of a total of 36,100 pixels, phase 3 occupies approximately 11,925 pixels of 33.03% of the total diagram. When an obstacle is added to the arena, phase 3 only occurs when *w_b_* = 50 and occupies approximately 4,500 pixels or 12.47% of the total diagram. Most of the parameter space that makes up phase 3 in the phase diagram for simulations without obstacles becomes phase 1 in the diagram with the obstacle; the remaining phase 3 region becomes phase 2 in the diagram with the obstacle. For both conditions, phase 3 never occurs when there is no directional bias (*w_b_* = 0) or when the concentration threshold is maximal (*T* = 1.0). Lastly, we compare the temporal dynamics of phase 3 simulations in Fig. 3C (time-series averaged over three repetitions). The average distance to the queen decreases faster and converges to a lower value when there is no obstacle (Fig. 5A). With the obstacle, we observe a slower decrease due to the exploration required to find the opening, and a convergence at a higher distance due to some bees that are still behind the obstacle by the end of the simulation. Similarly, the average growth in the queen’s cluster size over time is faster and the final size is larger when there is no obstacle (Fig. 5B). Both the average number of scenting bees (Fig. 5C) and average distance of the farthest scenting bee (Fig. 5D) follows similar trends of sharp initial increase and plateau (number of scenting bees) or slight decrease (farthest active distance), but there is a delay in the sharp increase when the obstacle is present.

**Figure 4:**
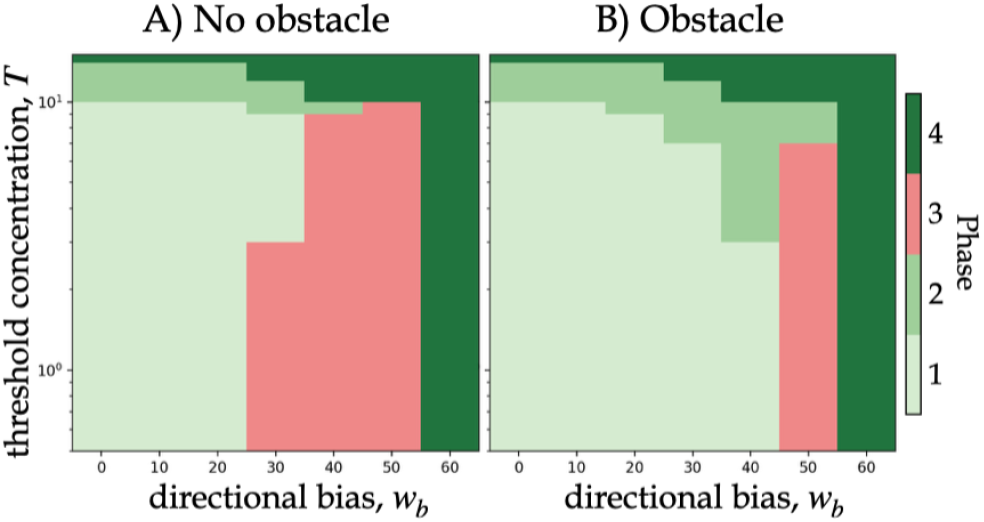
The effect of a physical obstacle on the phase boundaries for all simulations with varying *w_b_* and *T*. Phase diagrams constructed from scenting model dynamics using summary heatmaps of the final number of clusters, the final queen’s cluster size, and the distance of the farthest active bee to the queen. Phase 3 is highlighted in pink. A) Phase diagram without obstacle. B) Phase diagram with obstacle.

**Figure 5:**
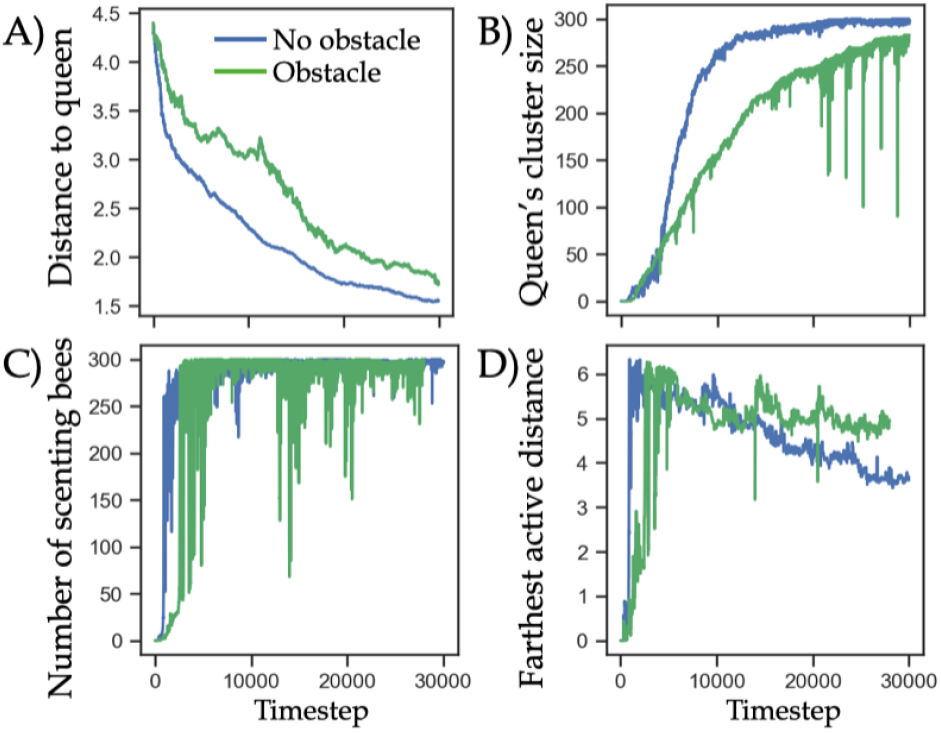
Time series of phase 3 simulations in Fig. 3C with and without obstacle (green and blue curves, respectively). A) The average distance of the worker bees to the queen over time. B) The average size of the queen’s cluster over time. C) The average number of scenting bees over time. D) The average distance of the farthest scenting bee over time.

## Discussion

Experimental studies show that bees employ the collective scenting strategy when localizing the queen and aggregating around her to form a coherent swarm (Nguyen et al., 2021b). This communication method is observed in a simple environment free of any perturbations as well as a more complex environment with the presence of physical obstacles (Nguyen et al., 2021b,a). The experimental results demonstrate a temporal delay in the peak of collective scenting and a slower decrease in the average bee distance to the queen as the obstacle requires the bees to first explore the space and find the opening before forming the scenting network.

While the experiments illuminate the temporal dynamics of localization in the presence and absence of obstacles, we also turn to modeling to better understand how the behavioral parameter space is affected by the environmental perturbation. The model predicts four distinct outcomes, or phases, determined by the directional bias and concentration threshold, (*w_b_*, *T*): 1) many small clusters, 2) clustering via only random walk, 3) clustering via signal propagation, and 4) no clustering. All four phases are present in both environments. However, the boundaries of the phases shift in the presence of the obstacle. Phase 4, dictated by very high *T* and very high *w_b_*, remains constant. However, the successful aggregation of phase 3 occupies a smaller overlapping area of the phase diagram with the obstacle, constraining the parameters to only one value of high *w_b_* = 50, while the phase spans *w_b_* values of 30, 40, and 50 in the diagram for simulations without the obstacle. With low values of *T*, most of the area encompassing phase 3 in the diagram without obstacles becomes phase 1 in the diagram with obstacles—the physical wall leads to the breaking of the chains of scenting bees in the percolation network, resulting in the formation of small local clusters on both sides of the obstacle. The phase 3 region with high values of *T* becomes phase 2 with the obstacle, as the wall separates bees from one another and prevents the percolation network from forming.

Nguyen et al. (2021b) shows the effect of bee density on phase boundaries: As density increases, there are more bees to create and sustain the communication network, and the range of *w_b_* and *T* and the area in the phase diagram for phase 3 is greater. In this study, the shrinking of phase 3 in the presence of the obstacle suggests that the wall has a similar effect to decreasing total density, by decreasing the effective density of scenting bees available to form a robust scenting percolation network. The bees are capable of navigating the obstacle environment to localize the queen and aggregate around her, but their range of potential behavioral parameters for the task is more limited and less flexible.

The model offers a simplified simulation of the bee collective scenting behavior. Some caveats and limitations that prevent the simulations from better matching the experimental data include the lack of spontaneous scenting observed in real bees but not modeled due to a lack of better understanding of the mechanisms. Further, bees in experiments seemingly stop scenting as they gather at the queen’s cluster, while we simply allow bees to continue scenting without a stop function. Future analysis of the experiments is required to understand the mechanisms of these behaviors in order to improve the accuracy of the model.

Additional future directions include testing various densities in simulations with obstacles to further understand the density effect on the phase diagram and the temporal dynamics of aggregation. There may be a critical density below which the successful aggregation via propagation in phase 3 will not be present, as the obstacle dramatically reduces the effective density of scenting bees. Further, the physical obstacle in this study is relatively simple; more complex physical obstacles or other kinds of environmental perturbations (e.g., varying opening size in the obstacle, multiple obstacles, a maze, wind, or a moving queen) are of interest and can further shed light on how bee swarms navigate the complex, noisy environments found in nature. Lastly, understanding how the behavioral parameters shift in varying environments for the bees may further inform the design and development of swarm robotic systems to include a parameter space that allows for successful group coordination in the presence of physical obstacles.

## Acknowledgements

This work was supported by the National Science Foundation Graduate Research Fellowship Program under Grant No. DGE1650115 (D.M.T.N.) and Physics of Living Systems Grant No. 2014212 (O.P.). Any opinion, findings, and conclusions or recommendations expressed in this material are those of the authors(s) and do not reflect the views of the NSF. We thank Chantal Nguyen for valuable feedback on this manuscript.

## Appendix

### Line-of-sight method

For a given line with end coordinates (*x*_1_, *y*_1_) and (*x*_2_, *y*_2_), its slope is *m* = (*y*_2_ − *y*_1_)/(*x*_2_ − *x*_1_) and y-intercept is *b* = −*mx*_1_ + *y*_1_. The equation of the line in standard form, *Ax* + *By* = *C*, is: *mx* − *y* = −*b*, where the constant *A* = *m, B* = −1, and *C* = −*b*. The system of equations for the 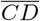 and 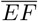 is: 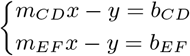

Transforming to a matrix equation to solve, where *A_CD_* = *m_CD_*, *A_EF_* = *m_EF_*, *B_CD_* = −1, *B_EF_* = −1, *C_CD_* = −*b_CD_*, 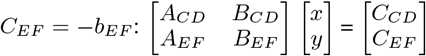

### Additional figures, table, and algorithm

**Figure A1:**
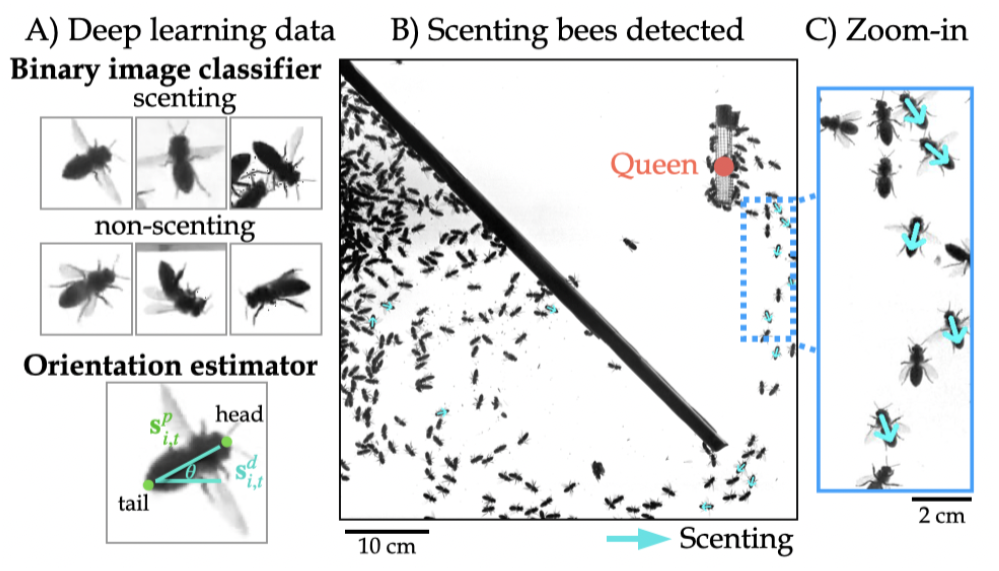
A) Training data examples for deep learning models to classify scenting bees and estimate orientations. B) Detections of scenting bees and their orientations (teal arrows). C) Zooming into scenting bees with wide wings.

**Figure A2:**
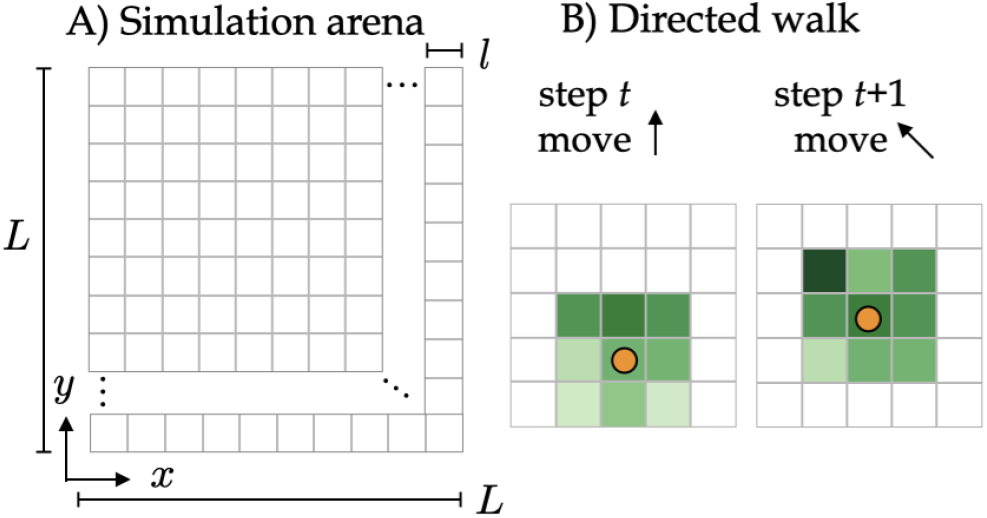
A) The *L* × *L* 2-d simulation arena is discretized into *l* × *l* sized pixels. B) When a bee detects a local pheromone concentration above the activation threshold (*C*(*x, y, t*) > *T*), it computes the gradient around it (using the nearest 9 pixels, highlighted in different shades of green) and walks up the gradient towards higher concentration.

#### Algorithm 1: Queen localization and aggregation algorithm

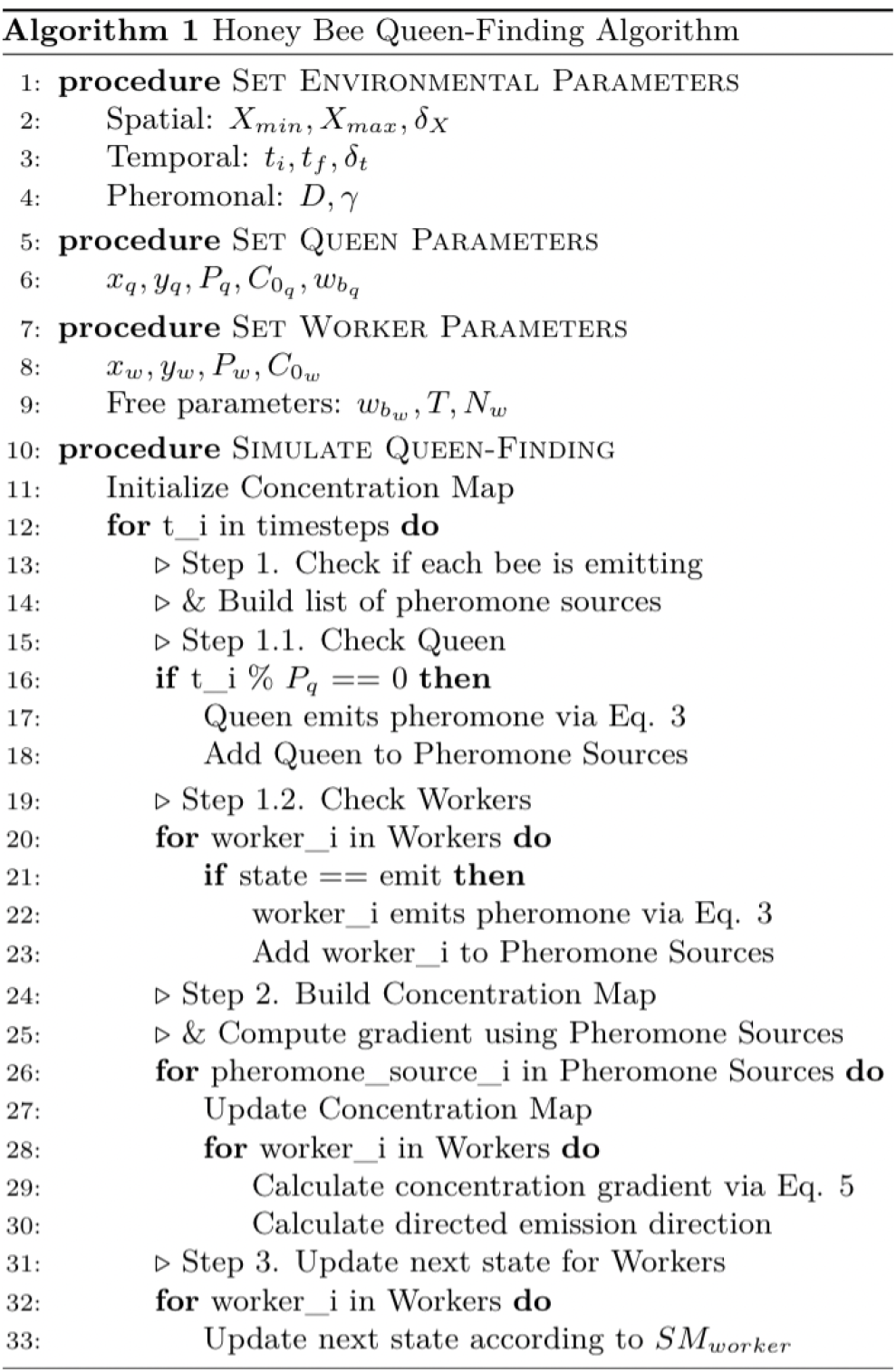

**Table 2:** Free parameters, *w_b_* and *T*, of the ABM are bolded.

| Parameter | Meaning | Value |
| --- | --- | --- |
| $X_{min}$ | Min size of grid arena | -3 |
| $X_{max}$ | Max size of grid arena | 3 |
| $\delta_X$ | Grid cell size | 0.01 |
| $t_i, t_f$ | Start and final time | 0, 150 |
| $\delta_t$ | Time integration constant | 0.005 |
| $D$ | Diffusion coefficient | 0.6 |
| $\gamma$ | Decay constant | 108 |
| $(x_q, y_q)$ | Queen 2-D position | (1,-2) |
| $P_q$ | Queen emission frequency | 80 |
| $C_{0q}$ | Queen initial concentration | 0.0575 |
| $w_{bq}$ | Queen bias scalar | 0 |
| $(x_w, y_w)$ | Worker 2-D position | $\in [X_{min}, X_{max}]$ |
| $P_w$ | Worker emission period | 80 |
| $C_{0w}$ | Worker initial concentration | 0.0575 |
| $N_w$ | Worker density | 300 |
| $w_{bw}$ | **Worker bias scalar** | [0, 10, 20, 30, 40, 50, 60] |
| $T$ | **Worker activation threshold** | [1e-4, 1e-3, 0.01, 0.025, 0.05, 0.075, 0.1, 0.20, 0.25, 0.3, 0.4, 0.5, 0.6, 0.7, 1.0] |

